# The biological sink of atmospheric H2 is more sensitive to spatial variation of microbial diversity than N_2_O and CO_2_ emissions in an agroecosystem

**DOI:** 10.1101/2021.10.15.464264

**Authors:** Xavier Baril, Audrey-Anne Durand, Narin Srei, Steve Lamothe, Caroline Provost, Christine Martineau, Claude Guertin, Kari Dunfield, Philippe Constant

## Abstract

The relationship between soil microbial diversity and agroecosystem functioning is controversial due to the elevated diversity level and the functional redundancy of microorganisms. A field trial was established to test the hypothesis that enhanced crop diversity with the integration of winter cover crops (WCC) in a conventional maize-soy rotation promotes microbial diversity and the biological sink of H_2_ in soil, while reducing N_2_O emissions to the atmosphere. *Vicia villosa* (hairy vetch), *Avena sativa* (oat), and *Raphanus sativus* (Daikon radish) were cultivated alone or in combinations and flux measurements were performed throughout two subsequent growing seasons. Soil acted as a net sink for H_2_ and as a net source for CO_2_ and N_2_O. CO_2_ flux was the most sensitive to WCC whereas a significant spatial variation was observed for H_2_ flux with soil uptake rates observed in the most productive area two-fold greater than the baseline level. Sequencing and quantification of taxonomic and functional genes were integrated to explain variation in trace gas fluxes with compositional changes in soil microbial communities. Fungal communities were the most sensitive to WCC, but neither community abundance nor beta diversity were found to be indicative of fluxes. The alpha diversity of taxonomic and functional genes, expressed as the number of effective species, was integrated into composite variables extracted from multivariate analyses. Only the composite variable computed with the inverse Simpson’s concentration index displayed a reproducible pattern throughout both growing seasons, with functional genes and bacterial 16S rRNA gene defining the two most contrasting gradients. The composite variable was decoupled from WCC treatment and explained 19-20% spatial variation of H_2_ fluxes. Sensitivity of the trace gas exchange process to soil properties at the local scale was inconsistent among H_2_, N_2_O and CO_2_, with the former being the most related to microbial diversity distribution pattern.

## Introduction

The agricultural sector contributes to nearly 12% global anthropogenic greenhouse gas (GHG) emissions (Smith et al., 2007). In temperate agroecosystems, the accelerated decomposition rate of soil organic matter (SOM) caused by tillage and the harvest of plant biomass contribute to the release of CO_2_ and application of excess fertilizers promotes N_2_O emissions. Land management practices reducing organic matter decomposition rates and increasing the input of C in soil are beneficial to mitigate GHG emissions and augment the CO_2_ sink capacity. As such, winter cover crops (WCC) are seen as a sustainable option to mitigate and adapt to climate change. WCC are integrated after the harvest of cash crops in organic and conventional crop rotations. WCC biomass residues are either removed or integrated in soil before the seedlings of subsequent cash crops. Their implementation as green manure is expected to cause soil organic carbon (SOC) accumulation at a rate of 1.2 Mg-CO_2_ equivalents ha^-1^ yr^-1^, independently from legume or non-legumes cover crop types, climate conditions and tillage or no-tillage management practices (Poeplau & Don, 2015). Furthermore, WCC were shown to improve soil fertility, enhance subsequent cash crop yield, improve soil structure, reduce soil erosion, control weeds and interrupt diseases and pest cycles (Abdallahi & N’dayegamiye, 2000; Cherr et al., 2006). This work examines two attributes of agroecosystems that are responsive to the integration of WCC in culture rotations: microbial diversity and trace gases fluxes.

WCC promotes soil belowground diversity but neither the intensity nor the resulting changes in functioning are constrained. The feedback response of microorganisms towards SOM inputs is likely the best documented mechanism relating WCC to microbial diversity. The incorporation of plant materials in soil activates a succession of microbial processes initiated with saprophytic guilds hydrolyzing lignocellulosic plant materials, mobilizing nutrients and participating in soil aggregates formation (Tiemann et al., 2015). A recent meta-analysis examining soil microbial diversity in 60 WCC field trials led to the conclusion that the implementation of WCC as green manure can lead to a modest increase of soil microbial diversity (Kim et al., 2020). The abundance and the activity of microbial communities appeared more responsive to WCC than microbial diversity, but environmental factors encompassing climate, tillage and soil types exerted confounding effects. Another potential factor deciphering the strength of the relationship between WCC and soil microbial diversity is the complexity and functional attributes of WCC. The coupling between aboveground and belowground diversity is governed by the plant-mediated fluxes of nutrients, energy and signaling molecules in soil. As the composition and rate of fluxes vary among different plant species, the diversity of aboveground biomass exerts a filtering effect on the structure and functioning of soil microbial communities (Philippot et al., 2013a). These interactions were mostly examined in grasslands, whereas supporting experimental evidence is lacking in crop rotations implemented in agroecosystems. In a short-term field trial for instance, management practices including tillage and glyphosate application were stronger drivers of soil microbial community structure than WCC mixtures comprising between two and eight WCC species (Romdhane et al., 2019). Examination of microbial abundance and soil enzyme activity in a field trial comprising up to eight WCC species led to similar observation after two successive growing seasons, providing no evidence supporting more benefits with WCC mixtures (Housman et al., 2021). More investigations are needed to examine the benefits of WCC mixture on soil microbial diversity and functioning, particularly for specialized functions involving rare members of microbial communities.

One of the microbial processes that have received attention in the context of crop rotations is the exchange of climate-relevant trace gases. Because WCC increases SOM, variations of gas exchanges are expected. Except for CO_2_, the response of gas fluxes in agriculture to WCC implementation is mostly site-specific. WCC tend to increase CO_2_ emission by the rise of SOC that they provide (Muhammad et al., 2019). The stability of the response among different biomes can be attributed to the broad diversity of microorganisms participating in CO_2_ emissions. Beside CO_2_, other greenhouse gases have been examined in crop rotations including WCC, with N_2_O being the most represented. Terrestrial ecosystems are the most important source of N_2_O, responsible for 70% of total emissions. Agricultural lands are accountable for almost half of these emissions (Syakila & Kroeze, 2011). N_2_O emissions are driven by microbial nitrification and denitrification processes in soil. The abundance and diversity of denitrifying bacteria determine the denitrification potential in soil, which is linked to N_2_O emission. (Philippot et al., 2011; Philippot et al., 2013b). The input of SOC in soil originating from WCC biomass and root exudates is favoring the growth of denitrifying communities, including specialized clades displaying a high affinity towards N_2_O, contributing to lesser net emissions to the atmosphere (Wang et al., 2021).

This response varies with the functional group of WCC, with non-legume WCC decreasing N_2_O emissions and legume WCC promoting emissions due to the opposite effect on the C/N ratio in soil (Basche et al., 2014; Muhammad et al., 2019). Less is known about the impact of WCC diversity on N_2_O emissions. The incorporation of plant residues from a mixture of 15 WCC in soil microcosms led to lower N_2_O emission than soil microcosm amended with vetch biomass residues alone (Drost et al., 2020). This response was specific for the vetch because neither the application of residues from radish nor oat alone promoted N_2_O emissions, supporting the observation of higher fluxes in legume WCC. Contrary to CO_2_ and N_2_O, nothing is known regarding the impact of WCC on H_2_ fluxes. H_2_ is an indirect GHG due to its reactivity towards hydroxyl radicals in the atmosphere. The biological sink of H_2_ is responsible for ∼80 % of the global losses of tropospheric H_2_ (Constant et al., 2009). This process involves high-affinity H_2_-oxidizing bacteria (HOB) represented by a broad diversity encompassing the phyla *Actinobacteria, Proteobacteria, Acidobacteria*, and *Chloroflexi* (Constant et al., 2011; Greening et al., 2016). Despite their ubiquitous distribution in soil, HOB functioning is sensitive to composition change of soil microbial communities (Saavedra-Lavoie et al., 2020). Whether the critical threshold of diversity required for the function is secured in the environment or not remains elusive, but other variables including soil moisture and carbon content are also important drivers of H_2_ uptake rates (Ehhalt & Rohrer, 2009). Enhancement of SOC triggered by WCC is therefore expected to promote the biological sink of atmospheric H_2_ in soil.

This study was aimed at examining the short-term response of trace gas fluxes and soil microbial communities after the implantation of three WCC species either alone or in combination. Enhancement of CO_2_ emissions and H_2_ soil uptake were expected in response to WCC treatments, whereas reduction of N_2_O emissions were anticipated in rotations comprising WCC. These responses were expected to be in line with microbial diversity and abundance, expressed upon the examination of bacterial, fungal, H_2_-oxidizing and denitrifying communities.

## Materials and Methods

### Field trial description and sampling

The work was conducted in a farm located at Saint-Joseph-du-Lac, Quebec, Canada (45.55 N 74.05 W). The site is managed under a conventional soybean-maize rotation with annual tillage. In 2018, an area of the farm was dedicated to the establishment of a field trial consisting in corn (*Zea mays L*.)-WCC-soybean rotations organized as a randomized complete block design with three replications for each WCC treatment (figure 1). Three WCC were selected according to soil hydrological properties, weak winter hardiness of plants and targeted ecosystem services using the Cover Crops Decision Tool (http://decision-tool.incovercrops.ca). The N_2_-fixing legume hairy vetch (*Vicia villosa*) was selected on the basis of the ammonium they supply to soil. Oat (*Avena sativa*) was expected to increase soil organic matter while suppressing both weed and soil compaction. Daikon radish (*Raphanus sativus*) produces natural chemical agents to suppress weeds, fungal pathogens and insects, in addition to developing approximately two feet taproots scavenging nitrogen from deep in the soil profile and increasing rooting depth opportunities for successive crops. The three species were either cultivated alone or in mixtures comprising two or three species, resulting in seven different combinations. Each replicated block comprised one unmanaged fallow control plot (7 WCC combinations + 1 control = 8 plots per block), for a total of 24 plots (11 m x 7.4 m). Treatments were assigned randomly and WCC were seeded after corn harvest in August 2018. Light tillage was applied in the spring to incorporate plant aboveground biomass and residues in soil.

**Figure 1.**
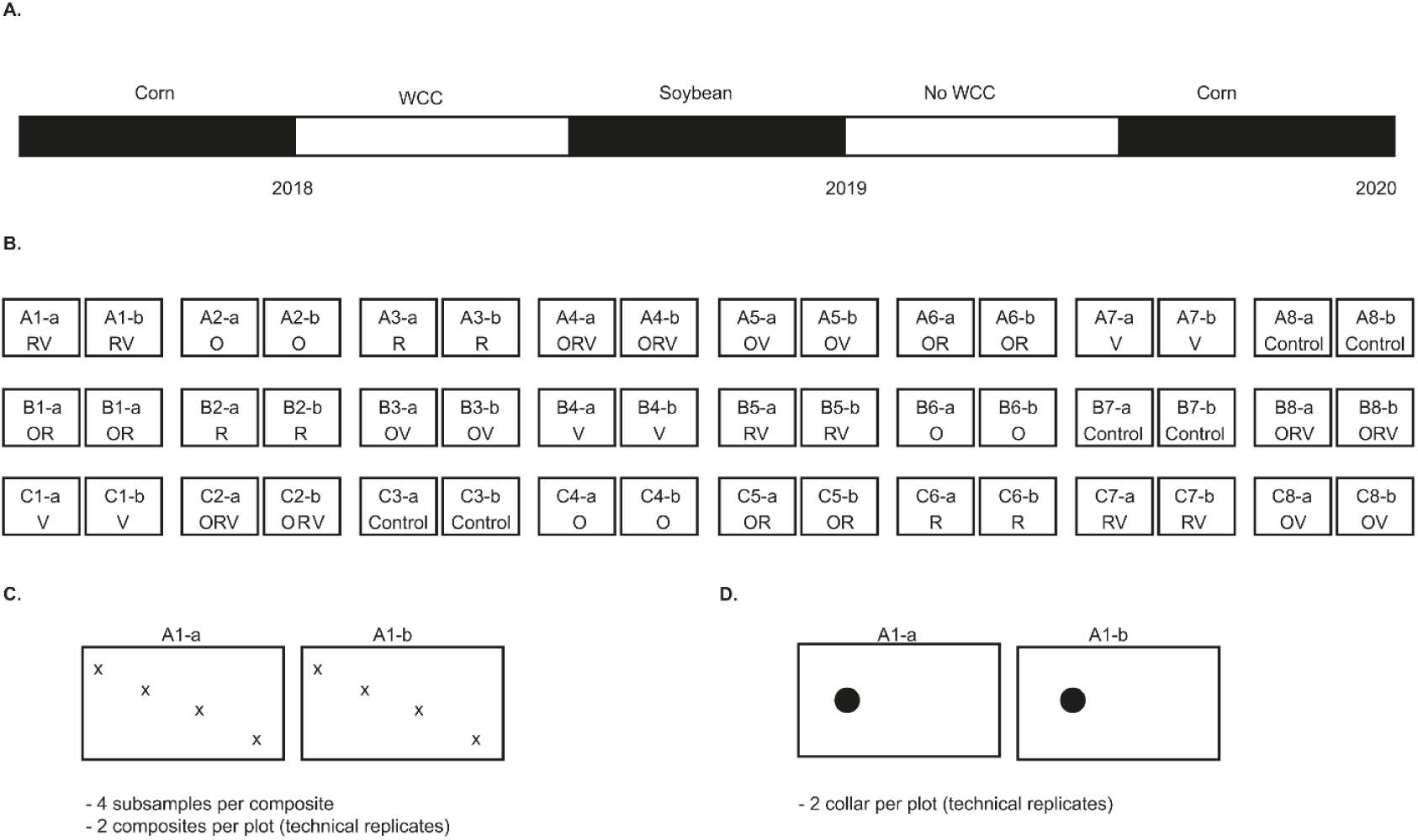
Schematic representation of the field trial. (a) The crop rotation sequence included a single application of WCC treatments in 2018. (b) Distribution of the WCC treatments was randomized in 24 plots distributed in three blocks. Each treatment is represented by two subplots designed by a capital letter (*A, B or C*) specifying block, a number (*1 to 8*) specifying the plot, and a suffix *(−a or -b*) specifying subplot. Each subplot is 4.5 m wild and 7.4 m long. Subplot are separated by 2 m and plot are separated by 5 m. O; Oat, R; Radish, V; Hairy Vetch. (c) A composite sample was assembled from each subplot by mixing four subsamples collected along a diagonal transect represented by the *x* characters. (d) A collar (black circle) was deployed at each subplot for flux measurements.

Each plot was separated into two subplots whose observations were taken as paired technical replicates (*i*.*e*., the average of values measured in paired technical replicates was computed prior to statistical analyses). Vegetation biomass was sampled from each subplot in October 2018 (WCC), October 2019 (soy) and August 2020 (corn) and dry weight was measured with a standard gravimetric method. Soil compaction was monitored in June 2018, June 2019 and July 2020 using a penetrometer (Field Scout SC-900®). Measurements were done at four locations along a diagonal transect in the subplots and readings were recorded at 0, 2.5, 5, 7.5 and 10 cm depth. Surface soil samples (0-5 cm) were collected in June 2018, 2019 and 2020. A composite sample was elaborated for each subplot with four subsamples collected along a diagonal transect (figure 1c). Genomic DNA was extracted from soil composite samples (0.25 g) with the DNeasy PowerLyzer PowerSoil kit (Qiagen®) and extracted DNA aliquot were stored at −20 °C. Composite samples were subjected to nutrients analysis performed at the Laurentian Forestry Centre (Sainte-Foy, Québec, Canada). Total C and N were obtained by combustion and gas chromatography (CNS928, LECO Corporation®, St. Joseph, USA). The Mehlich 3 extractant technique followed by measurements using the PerkinElmer® Optima™ 7300 DV ICP-OES instrument (PerkinElmer, Inc., Shelton USA) were utilized for P and K determinations as well as for other nutriment determination (Carter & Gregorich, 2007). Soil pH (1 g soil suspension in 10 ml 0.01 M CaCl_2_ solution) was measured with an accumet® model 15 pH-meter (FisherScientific, Pittsburgh, USA).

### Flux Measurements

CO_2_, H_2_ and N_2_O flux measurements were done with a static flux chamber technique. Each subplot comprised one permanent flux chamber collar (figure 1d). Collars (20 cm diameter) were installed one week before the beginning of flux measurement campaigns conducted from June 19^th^ to September 3^rd^ 2019 and June 16^th^ to August 11^th^ 2020. Barren soil was kept in the collar by removing debris and growing vegetation. Flux measurements were done two days a week, from 6h00 to 12h00. Sampling routine of the 24 plots was randomized with the function “sample” in the software R (R Core Team, 2013). A new sampling routine was defined each week, enabling a random survey of the subplots. Under this sampling strategy, the 24 plots were represented each week with measurements performed on a single collar. Flux measurements were done with a Licor analyzer control unit and 20 cm survey chamber model LI-8100 (LI-COR®, Lincoln, USA) operated with a septum placed in the air circulation network. The flux chamber was placed on the collar and 10 ml gas samples were collected every 2 minutes for a total of 10 minutes with a Pressure Lok® gastight glass syringe (VICI® Precision Sampling Inc., Baton Rouge, Louisiana, USA). Collected gas samples were transferred in conditioned Extainer®. Vials conditioning was done one day before sampling by performing five successive gas filling at 1.34 atm with 99.9999% N_2_ (Praxair) and vacuum cycles. Three blanks containing N_2_ were carried in the field and included in gas analyses routine to set the detection limit of the gas chromatographic assays. Gas samples were stored less than 30 h before analyzes. The background level of H_2_, CO_2_ and N_2_O was 237 ± 94 ppbv, 15.5 ± 8.9 ppmv and 0.009 ± 0.012 ppmv, respectively could be lowered with the method proposed by Nauer et al. (2021). A gas chromatography system equipped with a reductive gas detector (ta3000R, Ametek Process Instruments®, Delaware, USA) was utilized for H_2_ analyses and a SP1 7890-0504 gas chromatography system comprising a flame ionization detector and an electron capture detector (Agilent Technologies, CA, USA) was utilized for CO_2_ and N_2_O. A first reference gas mixture containing 2.0 ppmv H_2_ in air (Praxair, Morrisville, USA) and a second mixture containing 1 ppmv N_2_O and 600 ppmv CO_2_ in air (Greenhouse Gas Checkout Sample, Agilent Technologies Inc., Wilmington, USA) were used for calibration. All gas samples were processed for H_2_ measurements, while CO_2_ and N_2_O analyses were performed with samples collected at t = 0, 4, 8 and 10 min. Flux calculation and quality control were done with the package “Hmr” version 1.0.1 (Pedersen, 2020). Discrete flux measurements of the 24 plots over the course of two different days per week are prone to influences caused by sporadic rain or drought episodes. This variability on measured fluxes was smoothed by computing the median flux of each plot before testing the impact of WCC treatment or block on soil to air gaseous exchanges. Fluxes spatial variation along plots was generated with the package “rgeos” version 0.5.5 (Bivand et al., 2020).

### Abundance of Taxonomic and Functional Genes

The abundance of bacterial 16S rRNA gene, fungal ITS, *hhyL* gene encoding the large subunit of group 5/1h NiFe-hydrogenase, and *nosZ* gene encoding nitrous oxide reductase were determined by droplet digital PCR (ddPCR). Bacterial 16S rRNA was PCR-amplified with the primers Bakt_341F (5’-CCTACGGGNGGCWGCAG-3’) and Bakt_805R (5’-GACTACHVGGGTATCTAATCC-3’) (Herlemann et al., 2011), fungal internal transcribed spacer (ITS) with the primers ITS1F (5’-CTTGGTCATTTAGAGGAAGTAA-3’) and 58A2R (5’-CTGCGTTCTTCATCGAT-3’) (K. J. Martin & Rygiewicz, 2005), nitrous oxide reductase, *nosZ, nosZ*1-f (5’-WCSYTGTTCMTCGACAGCCAG-3’) and *nosZ*1-r (5’-ATGTCGATCARCTGVKCRTTYTC-3’) (Henry et al., 2006), [NiFe]-hydrogenase, *hhyL*, NiFe-244f (5’-GGGATCTGCGGGGACAACCA-3’) and NiFe-568r (5’-TCTCCCGGGTGTAGCGGCTC-3’) (Constant et al., 2011). The relative copy number concentrations of the subsample of 2019 and 2020 were measured by ddPCR Gene Expression EvaGreen® Assays (Bio-Rad, Hercules, USA). Genomic DNA extracted from the 48 composite soil samples was used for the ddPCR assays. Droplets were generated using the QX200 Droplet Generator (Bio-Rad). Sealed 96-well plate containing droplet emulsion was placed in a C1000 Touch™ Thermo Cycler (Bio-Rad). After PCR amplification (table S1), droplets were analyzed in a QX200™ Droplet Reader (Bio-Rad). The relative concentration for each subsample was obtained using QuantaSoft− software version 1.7.4 (Bio-Rad). In total, 8 runs of ddPCR were performed, with a random distribution of environmental DNA samples collected in 2019 and 2020 and comprising three negative controls with DNA-free sterile water. Manual threshold setting was used (table S2) and fixed to include rain in the positive fraction of the partitions (Huggett, 2020). Only samples with more than 10 000 accepted droplets were used for subsequent analyses. Assumed partition volumes were ∼20 000 nl (Bio-Rad Laboratories, 2015). Reported gene abundance is the average of the technical replicates represented by two composite samples collected in paired subplots.

### Diversity of Taxonomic and Functional Genes

Total genomic DNA was shipped to the *Centre d’expertise et de service Génome Québec* (Montréal, Québec, Canada) for PCR amplicon sequencing of bacterial 16S rRNA, fungal ITS region, *hhyL* gene and *nosZ* gene with the Illumina MiSeq PE-250 or MiSeq PE-300 platform. Raw sequence reads were deposited in the Sequence Read Archive of the National Center for Biotechnology Information under Bioproject PRJNA723322. Primer sequences were removed from raw reads with cutadapt (M. Martin, 2011). Downstream sequence quality control, amplicon sequence variant (ASV) clustering and taxonomic assignation was done with the package “dada2” version 1.16.0 (Callahan et al., 2015). Bacterial communities from 2018, 2019 and 2020 soil samples were characterized using 16S rRNA gene Amplicon Sequence Variant (ASV). For the four genes, a 100 reads cut-off was used to select the relevant ASVs for the analyses. Following this cut-off, 2702 ASV were kept for the 16S analyses. Fungal communities from 2019 and 2020 were characterized using ITS ASVs, where 632 ASVs were conserved following cut-off. Finally, after the cut-off, 389 and 1114 ASVs were kept for *hhyL* and *nosZ* analysis, respectively.

The taxonomic assignation of 16S rRNA gene and ITS region was based on SILVA version 138 (Quast et al., 2012) and UNITE general Fasta release version 10.10.2017 (UNITE Community, 2017) databases, respectively. Specificity of *hhyL* and *nosZ* ASV was verified against group 5/1h [NiFe]-hydrogenase database (Søndergaard et al., 2016), and *nosZ* sequences retrieved from the National Center for Biotechnology Information (NCBI) database (https://www.ncbi.nlm.nih.gov/) using the Basic Local Alignment Search Tool (Johnson et al., 2008). ASV sequences were converted to amino acid before their alignment against their corresponding reference database using Muscle (Edgar, 2004) implemented in the software MEGA-X version 10.2.4 (Kumar et al., 2018). Phylogenetic analysis of ASV and reference sequences was conducted with FastTree version 2.1.11 (Price et al., 2010). Alpha diversity of each gene was expressed with the first three Hill numbers representing species richness (q=0), the exponential function of the Shannon entropy index (q=1), and the inverse of Simpson index (q=2) (Hill, 1973). For each subplot, alpha diversity values were computed with the package “iNEXT” version 2.0.20 (Chao et al., 2014). The average of each paired subplot was computed to report alpha diversity value in the 24 plots.

### Statistical Analysis

Statistical analyses were performed using the software R version 4.0.4 (R Core Team, 2013). The effect of WCC treatments crop yield, soil physicochemical properties, trace gas fluxes, alpha diversity of microbial communities was examined with one-way ANOVA with the block and WCC treatment followed by a Tukey *post hoc* test with the package “stats” version 4.0.4 (R Core Team, 2013). Genes displaying the most important contribution in distinguishing alpha diversity amongst plots were identified with principal component analysis (PCA). One PCA was computed per alpha diversity metric (3 metrics) per year (2 years), resulting in six PCA. ITS region 16S rRNA, *hhyL* and *nosZ* gene alpha diversity variables were centered-reduced before the ordination of the 24 plots in the reduced space of PCA generated with the “princomp” function in the package “stats” version 4.0.4 (R Core Team, 2013). PCA plots were generated with the package “ggplot2” version 3.3.3 (Wickham, 2016). Coordinates of the 24 plots in the reduced space of the six PCA were used to elaborate composite alpha diversity indexes. The correlations between traces gas fluxes, diversity and soil physicochemical properties were examined with a Spearman’s rank correlation test computed with the package “stats” version 4.0.4 (R Core Team, 2013) and graphical outputs were generated with the package “corrplot” version 0.90 (Wei & Simko, 2021). Models to explain the variation of gas fluxes have been performed with simple or multiple linear regressions computed with the package “stats” version 4.0.4 (R Core Team, 2013) and graphical outputs were produced with the package “ggplot2” version 3.3.3 (Wickham, 2016). The effect of WCC treatment on beta diversity of soil microbial communities was tested with a Permutational multivariate analysis of variance (PERMANOVA), generated with the package “vegan” version 2.5.7 (Anderson, 2014; Jari Oksanen, 2020), to relate pairwise dissimilarity of microbial community profile to WCC treatments. Principal Coordinates Analysis (PCoA) was generated with the package “phyloseq” version 1.32.0 (McMurdie & Holmes, 2013) and graphical outputs were obtained with the package “ggplot2” 3.3.3 (Wickham, 2016) to visualise the similarities between community composition along plots.

Analysis of the phylogenetic diversity of the microbial community between treatments was computed with the package “picante” version 1.8.2 (Kembel et al., 2010), where the relationship between dissimilarity measures and treatments was obtained using a permutational MANOVA (AMOVA) test generated with the package “vegan” version 2.5.7 (Jari Oksanen, 2020). Distance-based redundancy analysis (db-RDA) were computed to relate the composition of microbial communities to trace gases fluxes and soil compaction with the package “vegan” version 2.5.7 (Jari Oksanen, 2020). Dispersion of microbial community structures from replicated WCC treatments was evaluated with Betadisper tests with the package “vegan” version 2.5.7 (Anderson, 2006; Jari Oksanen, 2020).

## Results

### Crop biomass, soil properties and trace gas fluxes

Aboveground biomass was harvested and partitioned to measure the dry weight of weed and WCC (figure 2). Biomass was undistinguishable amongst the different WCC treatment, except the control. Biomass was in the range 0.8 and 598 g_(dw)_ m^-2^ with weed representing between 0 and 39 g_(dw)_ m^-2^. Proportion of oat and radish were even in O+R mixtures, oat achieved higher biomass than hairy vetch in O+HV mixture and the biomass of Daikon radish was dominant in O+R+HV treatment. The sole WCC that survived winter Hairy vetch was incorporated in soil before seedlings in June 2019. The legacy effect of WCC was followed during the two subsequent growing seasons of soybean (2019) and maize (2020). WCC treatments did not explain variation of nutrient concentration, pH, cation exchange capacity, soil compaction or cash crop biomass amongst plots (ANOVA, *P* > 0.1). A block effect was noticed to explain variation of soil compaction (ANOVA, *P* < 0.05). In 2019, plots encompassing the block A (288 ± 25 kPa) were more compacted than those in block B (228 ± 37 kPa) and C (200 ± 33 kPa). A different pattern was observed in 2020 with the block B (1345 ± 256 kPa) showing higher soil compaction than block C (968 ± 267 kPa). In 2020, there was a block effect for C.E.C (ANOVA, *P* < 0.04). Bock C (0.13 ± 0.02 cmol(+) kg^-1^) had a higher content of C.E.C than block A (0.11 ± 0.01 cmol(+) kg^-1^).

**Figure 2.**
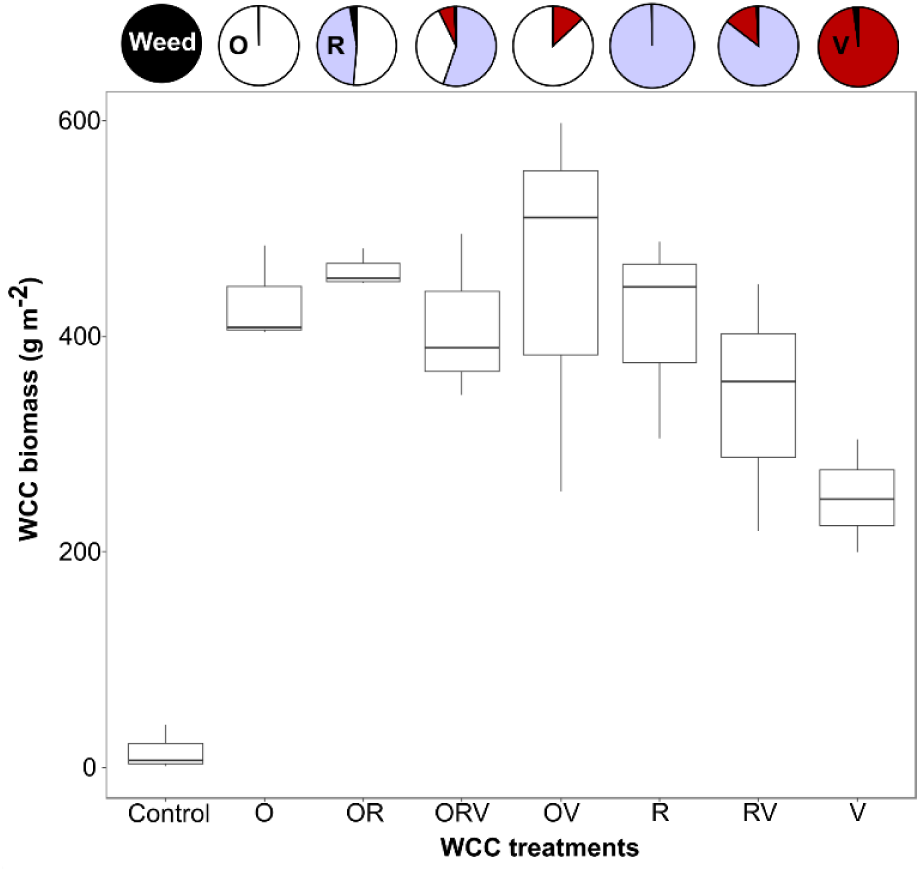
Biomass of WCC and their proportion in mixtures. The boxplot represents the dry weight of aboveground plant biomass in each WCC treatment. Pie charts of the upper panel show the proportion of WCC and weed. O; Oat, R; Radish, V; Hairy Vetch.

Fluxes were measured on a weekly basis during both growing seasons (figure 3). As a whole, soil acted as a net sink for H_2_ and as a net source for CO_2_ and N_2_O, with median fluxes in the range of -43 ± 31 µg m^-2^ h^-1^, 126 ±114 mg m^-2^ h^-1^, and 7 ± 11360 µg m^-2^ h^-1^, respectively (table 1). No block effect explained variation of trace gases fluxes (ANOVA, *P* > 0.05). H_2_, CO_2_ and N_2_O soil to air exchanges relying on different biological processes, the pattern of their monthly flux variations differed (figure 3). H_2_ soil uptake activity was the most stable process albeit a trend toward higher rates was observed in July and August of both growing seasons. The lowest CO_2_ fluxes were measured in June 2020, the month with the lowest total precipitation. In contrast, the higher N_2_O flux observed in the spring of the first growing season is consistent with total rain precipitations of 115 and 65 mm observed in June 2019 and 2020, respectively. Variation of trace gas fluxes was not explained by WCC treatments (ANOVA, *P* > 0.1). Nevertheless, a potential legacy effect of WCC was noticed for CO_2_ fluxes and H_2_ uptake rate that increased with WCC biomass in 2019 (ρ > 0.25, *P* < 0.005).

**TABLE 1:** Average values of gas fluxes and soil proprieties for each treatment in 2019 and 2020.

| Year | Treatment | H<sub>2</sub> flux<br>($\mu\text{g m}^{-2} \text{h}^{-1}$) | CO<sub>2</sub> flux ($\text{mg m}^{-2} \text{h}^{-1}$) | N<sub>2</sub>O flux<br>($\mu\text{g m}^{-2} \text{h}^{-1}$) | Soil compaction<br>($\text{kPa cm}^{-1}$) | Soil pH | Total C (%) | Total N (%) | C.E.C. (cmol(+) $\text{kg}^{-1}$) |
| --- | --- | --- | --- | --- | --- | --- | --- | --- | --- |
| 2019 | Control | 38 ± 11 | 167 ± 117 | 11 ± 15 | 223 ± 54 | 5.7 ± 0.1 | 3.3 ± 0 | 0.28 ± 0.01 | 33 ± 1 |
|  | O | 40 ± 7 | 175 ± 59 | 22 ± 21 | 227 ± 56 | 5.5 ± 0.2 | 3 ± 0.5 | 0.26 ± 0.03 | 33 ± 1 |
|  | OR | 32 ± 10 | 137 ± 26 | 7 ± 7 | 241 ± 29 | 5.8 ± 0.2 | 3.3 ± 0.2 | 0.3 ± 0.01 | 34 ± 3 |
|  | ORV | 52 ± 11 | 190 ± 25 | 21 ± 6 | 226 ± 68 | 5.7 ± 0.2 | 3.6 ± 0.4 | 0.31 ± 0.04 | 35 ± 1 |
|  | OV | 48 ± 7 | 210 ± 125 | 11 ± 5 | 269 ± 57 | 5.4 ± 0.3 | 3.5 ± 0.1 | 0.29 ± 0.01 | 33 ± 3 |
|  | R | 53 ± 25 | 188 ± 31 | 35 ± 36 | 221 ± 77 | 5.7 ± 0.4 | 3.5 ± 0.2 | 0.31 ± 0.02 | 33 ± 3 |
|  | RV | 41 ± 10 | 130 ± 71 | 8 ± 8 | 232 ± 26 | 5.9 ± 0.7 | 2.8 ± 0.6 | 0.24 ± 0.05 | 36 ± 4 |
|  | V | 36 ± 2 | 172 ± 24 | 12 ± 6 | 271 ± 39 | 5.4 ± 0.3 | 3.6 ± 0.4 | 0.32 ± 0.05 | 34 ± 4 |
| 2020 | Control | 48 ± 15 | 62 ± 22 | 6 ± 5 | 1201 ± 118 | 5.5 ± 0 | 3.4 ± 0.1 | 0.29 ± 0.02 | 31 ± 1 |
|  | O | 39 ± 3 | 63 ± 20 | 7 ± 2 | 1089 ± 169 | 5.7 ± 0.3 | 3.1 ± 0.4 | 0.28 ± 0.02 | 31 ± 2 |
|  | OR | 48 ± 6 | 117 ± 57 | 9 ± 7 | 1106 ± 311 | 5.6 ± 0.2 | 3.4 ± 0.3 | 0.31 ± 0 | 32 ± 1 |
|  | ORV | 47 ± 5 | 78 ± 19 | 7 ± 3 | 1206 ± 193 | 5.7 ± 0.1 | 3.6 ± 0.2 | 0.32 ± 0.03 | 32 ± 1 |
|  | OV | 39 ± 5 | 79 ± 57 | 10 ± 2 | 1104 ± 410 | 5.6 ± 0 | 3.4 ± 0 | 0.31 ± 0.01 | 31 ± 1 |
|  | R | 45 ± 2 | 142 ± 76 | 15 ± 15 | 1245 ± 51 | 5.7 ± 0.1 | 3.7 ± 0.2 | 0.33 ± 0.02 | 32 ± 0 |
|  | RV | 52 ± 13 | 70 ± 35 | 21 ± 28 | 1207 ± 413 | 5.8 ± 0.3 | 3 ± 0.6 | 0.26 ± 0.05 | 30 ± 1 |
|  | V | 47 ± 18 | 82 ± 26 | 4 ± 4 | 1047 ± 556 | 5.6 ± 0.2 | 3.6 ± 0.6 | 0.33 ± 0.06 | 33 ± 6 |

C, Carbon; N, Nitrogen; C.E.C, soil cation exchange capacity

**Figure 3.**
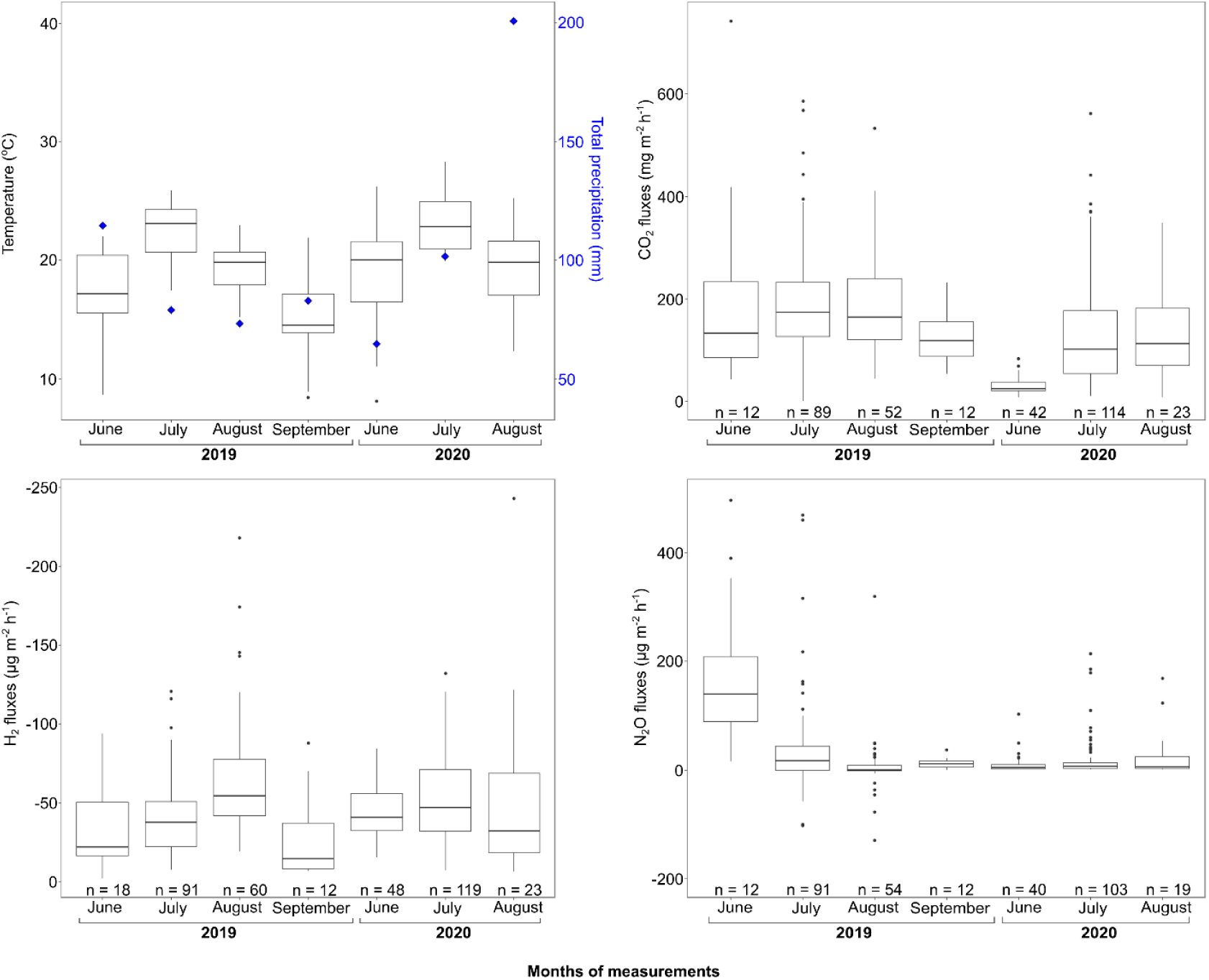
Monthly variation of meteorological factors and gas fluxes. The boxplot represents the average daily temperature or the measured gas fluxes in each month. The blue diamonds on the upper left panel represent the total rain precipitation for the month. For N_2_O fluxes, three measurements (> 600 µg N_2_O m^-2^ h^-1^) were not included to facilitate graphical visualization, these observations were made in July 2019 and in June and July 2020.

The biological sink of atmospheric H_2_ was the process displaying the largest spatial variation (figure 4). The intensity of H_2_ fluxes was irregular, with the most productive area displaying net uptake rates two-fold greater than the baseline level. In 2019, the five plots displaying the highest H_2_ uptake rates displayed an average flux of -61 ± 7 µg m^-2^ h^-1^ when compared to -38 ± 8 µg m^-2^ h^-1^ for the 19 other plots. In 2020, the highest H_2_ soil uptake rates were observed in three plots (−64 ± 2 µg m^-2^ h^-1^) compared to the baseline -43 ± 6 µg m^-2^ h^-1^ measured in the other plots. The position of plots exerting disproportionate H_2_ uptake rates was inconsistent between the two growing seasons, further supporting the negligible incidence of WCC treatment on H_2_ fluxes. CO_2_ and N_2_O did not show such a spatial distribution pattern.

**Figure 4.**
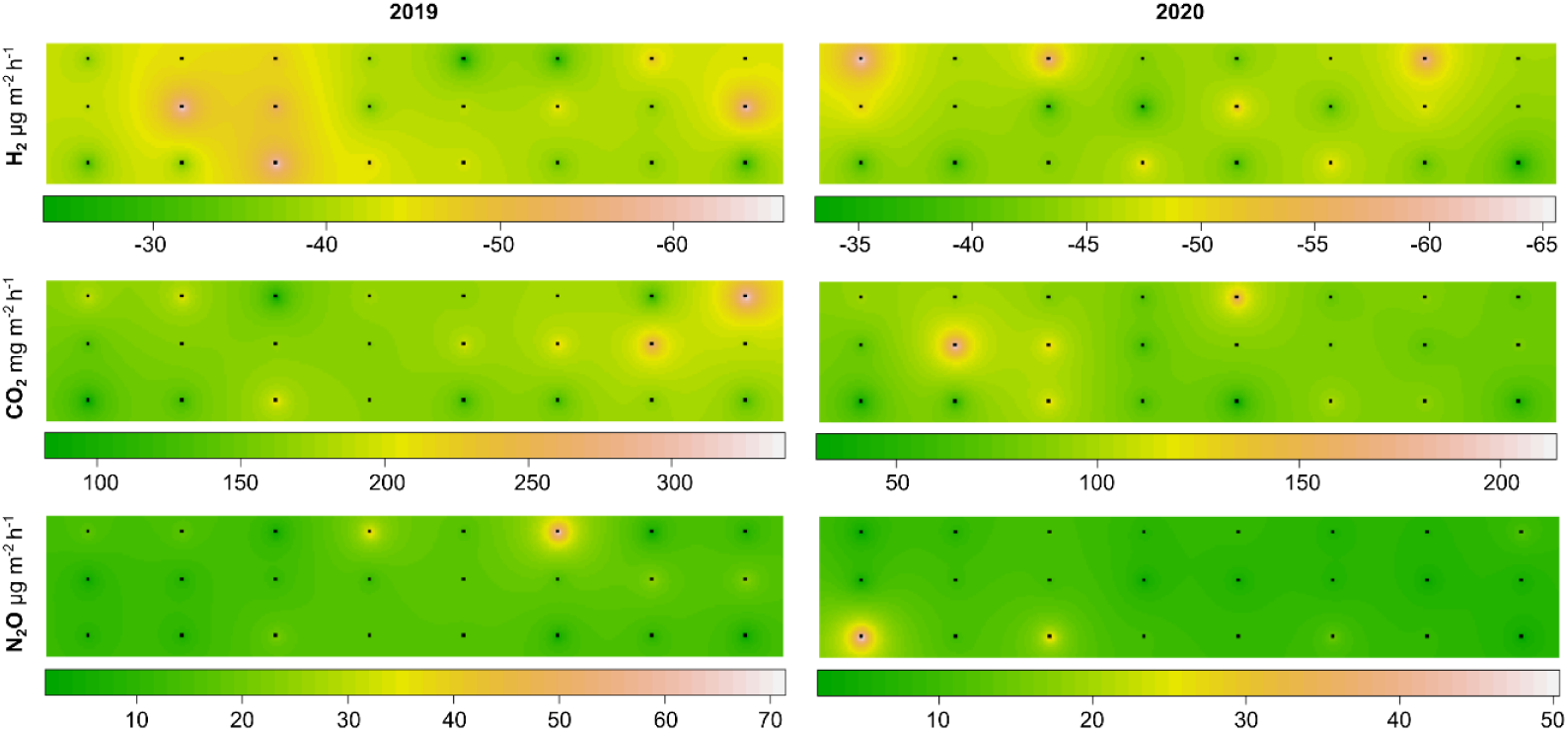
Spatial variation of gas fluxes along the plots. Color scale represents the median of gas fluxes for each plot represented by a black dot.

### Soil microbial communities

Diversity and abundance of bacterial 16S rRNA gene, fungal ITS, HOB *hhyL* gene and denitrifying bacteria *nosZ* gene were analyzed in the field trial. Alpha diversity expressed as species richness, Shannon entropy index and the inverse of Simpson index were greater for the 16S gene than for the other genes (table 2). Comparison of alpha diversity metrics of HOB and denitrifying bacteria provided new insights into the structure of those functional guilds that have never been simultaneously examined in the same ecosystem. The community of denitrifying bacteria displayed higher species richness and evenness than HOB. Although gene detection often is decoupled from functioning, this is a first evidence supporting potential higher resilience or resistance of the former group to disturbance than the second in the field trial. Neither the variation of alpha diversity metrics (ANOVA, *P*> 0.1) nor the variation of the abundance (ANOVA, *P* > 0.1) of taxonomic and functional gene markers was explained by WCC treatments. Simultaneous examination of taxonomic and functional groups was an opportunity to examine which component of microbial communities displayed the most important variation of alpha diversity in the experimental trial. This was achieved by using unconstrained multivariate analysis of the four marker genes (figure 5). The two first components of PCA explained 69 to 77% variation of alpha diversity distribution patterns, whereas the contribution of each gene marker in defining reduced spaces were not conserved amongst the three diversity metrics. Consistence of alpha diversity gradients amongst the two years appeared to increase with the Hill number of alpha diversity metrics. For instance, the second component of species richness was mainly driven by bacterial 16S rRNA gene in both growing seasons, but the corresponding first components were ITS and *nosZ* in 2019 and 2020, respectively. A more consistent ordination structure was observed for evenness of microbial communities. HOB communities were the most important driver of inverse of Simpson variation amongst the plots, followed by bacterial 16S rRNA gene for both growing seasons. Positions of plots along the two axes of PCA were integrated as two composite metrics (PCA1 and PCA2) representing the most prominent gradients of alpha diversity in soil of the field trial. None of the composite variables computed on the basis of either species richness, the Shannon entropy index or the inverse of Simpson index was related to WCC treatments (ANOVA, p > 0.1). Correlations were found between alpha diversity composite metrics and soil and plant variables including pH, compaction and WCC biomass. Their patterns were, however, inconsistent among the six composite metrics, impairing a sound interpretation of environmental features related to alpha diversity extremes at the field trial level.

**TABLE 2 :** Diversity indexes of taxonomic and functional genes for each treatment in 2019 and 2020.

| Year | Treatment | *hhyL* |  |  | *nosZ* |  |  | 16S rRNA gene |  |  | Fungal ITS |  |  |
| --- | --- | --- | --- | --- | --- | --- | --- | --- | --- | --- | --- | --- | --- |
|  |  | Species richness | Shannon index | Simpson index | Species richness | Shannon index | Simpson index | Species richness | Shannon index | Simpson index | Species richness | Shannon index | Simpson index |
| 2019 | Control | 188 ± 8 | 64 ± 5 | 30 ± 3 | 325 ± 27 | 95 ± 17 | 43 ± 10 | 948 ± 54 | 564 ± 59 | 342 ± 54 | 116 ± 9 | 40 ± 2 | 20 ± 1 |
|  | O | 179 ± 18 | 65 ± 8 | 32 ± 6 | 341 ± 18 | 106 ± 15 | 46 ± 13 | 950 ± 45 | 578 ± 37 | 353 ± 29 | 120 ± 20 | 38 ± 15 | 19 ± 9 |
|  | OR | 171 ± 19 | 55 ± 7 | 25 ± 5 | 362 ± 11 | 102 ± 23 | 40 ± 18 | 1015 ± 123 | 626 ± 69 | 375 ± 26 | 130 ± 9 | 40 ± 3 | 18 ± 2 |
|  | ORV | 174 ± 29 | 55 ± 15 | 24 ± 8 | 343 ± 6 | 89 ± 16 | 32 ± 13 | 997 ± 59 | 624 ± 36 | 405 ± 40 | 127 ± 5 | 34 ± 10 | 16 ± 7 |
|  | OV | 180 ± 8 | 59 ± 8 | 27 ± 5 | 346 ± 21 | 96 ± 21 | 39 ± 18 | 1023 ± 113 | 635 ± 80 | 406 ± 46 | 135 ± 5 | 43 ± 5 | 21 ± 3 |
|  | R | 164 ± 12 | 52 ± 9 | 24 ± 6 | 342 ± 9 | 107 ± 2 | 48 ± 1 | 1010 ± 24 | 639 ± 24 | 411 ± 33 | 126 ± 2 | 37 ± 5 | 18 ± 4 |
|  | RV | 168 ± 30 | 59 ± 17 | 28 ± 10 | 354 ± 4 | 110 ± 1 | 47 ± 4 | 1045 ± 185 | 651 ± 103 | 414 ± 56 | 134 ± 4 | 40 ± 11 | 19 ± 7 |
|  | V | 178 ± 38 | 61 ± 22 | 29 ± 13 | 338 ± 30 | 92 ± 18 | 36 ± 14 | 1058 ± 51 | 644 ± 48 | 378 ± 93 | 133 ± 17 | 40 ± 12 | 18 ± 9 |
| 2020 | Control | 100 ± 2 | 38 ± 1 | 20 ± 1 | 398 ± 21 | 138 ± 6 | 61 ± 6 | 568 ± 93 | 406 ± 64 | 289 ± 42 | 204 ± 16 | 74 ± 3 | 38 ± 1 |
|  | O | 94 ± 4 | 39 ± 3 | 23 ± 3 | 412 ± 20 | 134 ± 19 | 54 ± 16 | 535 ± 62 | 384 ± 35 | 268 ± 14 | 211 ± 14 | 75 ± 7 | 36 ± 4 |
|  | OR | 101 ± 6 | 41 ± 6 | 24 ± 5 | 398 ± 63 | 115 ± 45 | 44 ± 26 | 566 ± 109 | 404 ± 55 | 287 ± 21 | 213 ± 15 | 83 ± 6 | 45 ± 4 |
|  | ORV | 102 ± 15 | 35 ± 5 | 18 ± 3 | 401 ± 17 | 111 ± 22 | 40 ± 16 | 508 ± 81 | 369 ± 45 | 272 ± 21 | 231 ± 8 | 86 ± 5 | 47 ± 2 |
|  | OV | 106 ± 6 | 42 ± 4 | 23 ± 3 | 446 ± 23 | 142 ± 17 | 57 ± 12 | 517 ± 142 | 375 ± 92 | 264 ± 62 | 236 ± 17 | 84 ± 17 | 43 ± 13 |
|  | R | 99 ± 15 | 39 ± 7 | 22 ± 4 | 416 ± 36 | 125 ± 20 | 46 ± 14 | 575 ± 112 | 411 ± 65 | 293 ± 25 | 216 ± 6 | 75 ± 3 | 38 ± 4 |
|  | RV | 103 ± 9 | 41 ± 7 | 24 ± 5 | 388 ± 92 | 108 ± 17 | 36 ± 11 | 484 ± 190 | 347 ± 113 | 248 ± 61 | 204 ± 37 | 67 ± 24 | 35 ± 14 |
|  | V | 105 ± 10 | 41 ± 7 | 23 ± 6 | 384 ± 46 | 112 ± 44 | 43 ± 27 | 565 ± 48 | 408 ± 27 | 295 ± 22 | 211 ± 18 | 73 ± 8 | 38 ± 3 |

**Figure 5.**
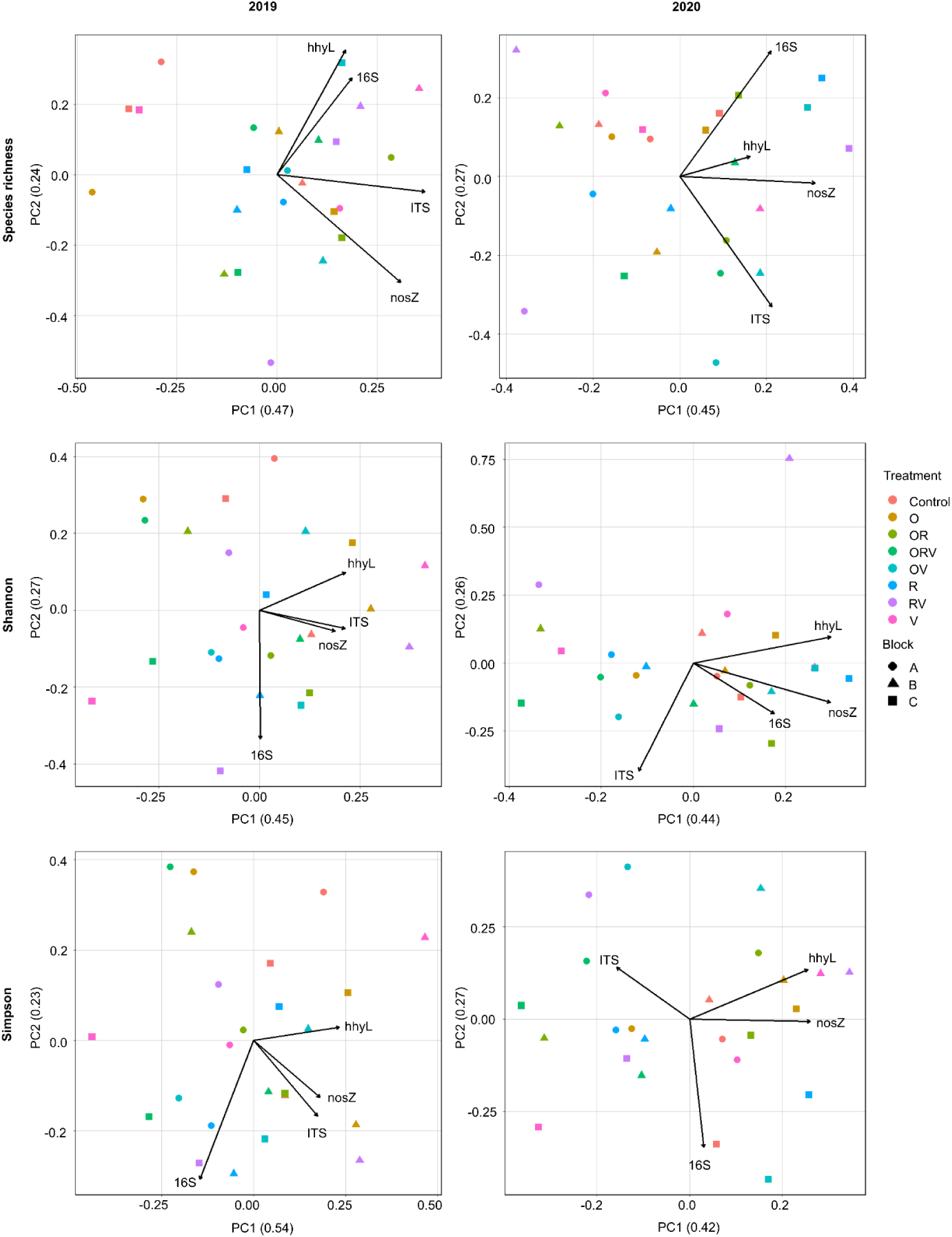
Principal Component Analysis (PCA) of plot alpha diversity indexes associated with each gene for the first three Hill numbers (q=0, 1, 2) in 2019 and 2020.

Composition of microbial communities was dominated by ASV encompassing the bacterial classes Alphaproteobacteria (32%) and Actinobacteria (15%) whereas fungi were dominated by Sordariomycetes (7%). Taxonomic assignation of HOB and denitrifying bacteria was not intended owing to frequent lateral transfer of functional genes encoding hydrogenase and nitrous oxide reductase (supplementary file), but the ASV1 (86% amino acid sequence identity to its closest relative *Schlesneria paludicola*) dominated HOB (10%) and the ASV1 displaying 75% amino acid sequence identity to *Neptunomonas qingdaonensis* dominated denitrifer communities (9%). The most important factor explaining variation of microbial composition was the sampling date for the four marker genes (Permanova, R^2^ > 0.25, *P* < 0.001). The overall pattern of bacterial communities was neither explained by WCC treatment nor experimental block (Permanova, R^2^ < 0.26, *P* > 0.06). For fungal communities, a block effect was observed in 2019 (R^2^ = 0.12, *P* < 0.004) but a potential legacy of WCC treatment was noticed in 2020. This effect was driven by HV and V treatments displaying different composition (Permanova, R^2^ = 0.35, *P* < 0.02) and heterogeneity with more homogeneous and less diverse ITS profiles for the V treatment (Betadisper, *P* < 0.05). These differences were not accompanied with microbial phylogenetic dissimilarity between plots (AMOVA, *P* > 0.05), indicating uneven instead of systematic covariations of taxonomic groups.

### Covariation of trace gas flux with microbial diversity and soil physicochemical attributes

Soil physicochemical parameters and microbial diversity patterns were integrated into a series of correlation and multivariate analyses to relate their variation to trace gases flux (figure S1). A few variables displayed consistent covariation with trace gases flux between the two growing seasons. CO_2_ fluxes were correlated with total carbon and total nitrogen (ρ > 0.30, *P* < 0.001). None of the measured soil or vegetation biomass variables displayed consistent correlation with N_2_O or H_2_ fluxes during both growing seasons. Species richness, Shannon entropy index and the inverse of Simpson index were examined for each marker gene alone or as composite metrics (PCA1 and PCA2). The bacterial Simpson index (ρ > 0.18, *P* < 0.002) and the Simpson index composite metric PCA2 (ρ > 0.22, *P* < 0.004) were correlated with CO_2_ fluxes. N_2_O fluxes were rather correlated with bacterial species richness (ρ > 0.11, *P* < 0.02). However, N_2_O fluxes also correlated with Shannon PCA1 (ρ > 0.03, *P* < 0.04). Both functional and taxonomic diversity were correlated with H_2_ fluxes, i.e., HOB Shannon entropy index and the inverse of Simpson index, and bacterial Simpson index (ρ > 0.12, *P* < 0.03). Thus, both PCA1 and PCA2 of the Simpson index correlated with H_2_ fluxes (ρ > 0.18, *P* < 0.02), where a stronger correlation was observed with PCA2 (ρ > 0.35, *P* < 0.001). No significant relationship was observed between the composition of taxonomic and functional communities and trace gas fluxes, according to distance-based redundancy analyses (data not shown).

Stepwise regression analyses were conducted to identify the factors contributing to the most important fraction of variation (figure 6). For CO_2_ flux, the single linear regression including total carbon as independent variable was the most parsimonious model for both growing seasons. The inverse of Simpson index composite captured CO_2_ fluxes variation in 2019 (R^2^ = 0.21, *P* < 0.03), but failed to explain 2020 CO_2_ fluxes variation, which was best explained by the biomass of WCC. Soil physicochemical variables were those capturing the most important variation of N_2_O fluxes, albeit no consistent response was observed for both growing seasons. Single linear regressions including either soil compaction in 2019 (R^2^ = 0.32, *P* < 0.004) or pH in 2020 (R^2^ = 0.30, *P* < 0.006) were optimal. For H_2_, the alpha diversity composite variable based on the inverse of Simpson index coordinates explained 19-20% (*P* < 0.04) flux variation. This relationship implies superior performance of the biological sink of atmospheric H_2_ in plots displaying the highest evenness within the community and a decoupling of the process with alpha diversity of HOB. Model resolution was enhanced by the addition of total soil carbon as second independent variable (R^2^ = 0.26, P < 0.02).

**Figure 6.**
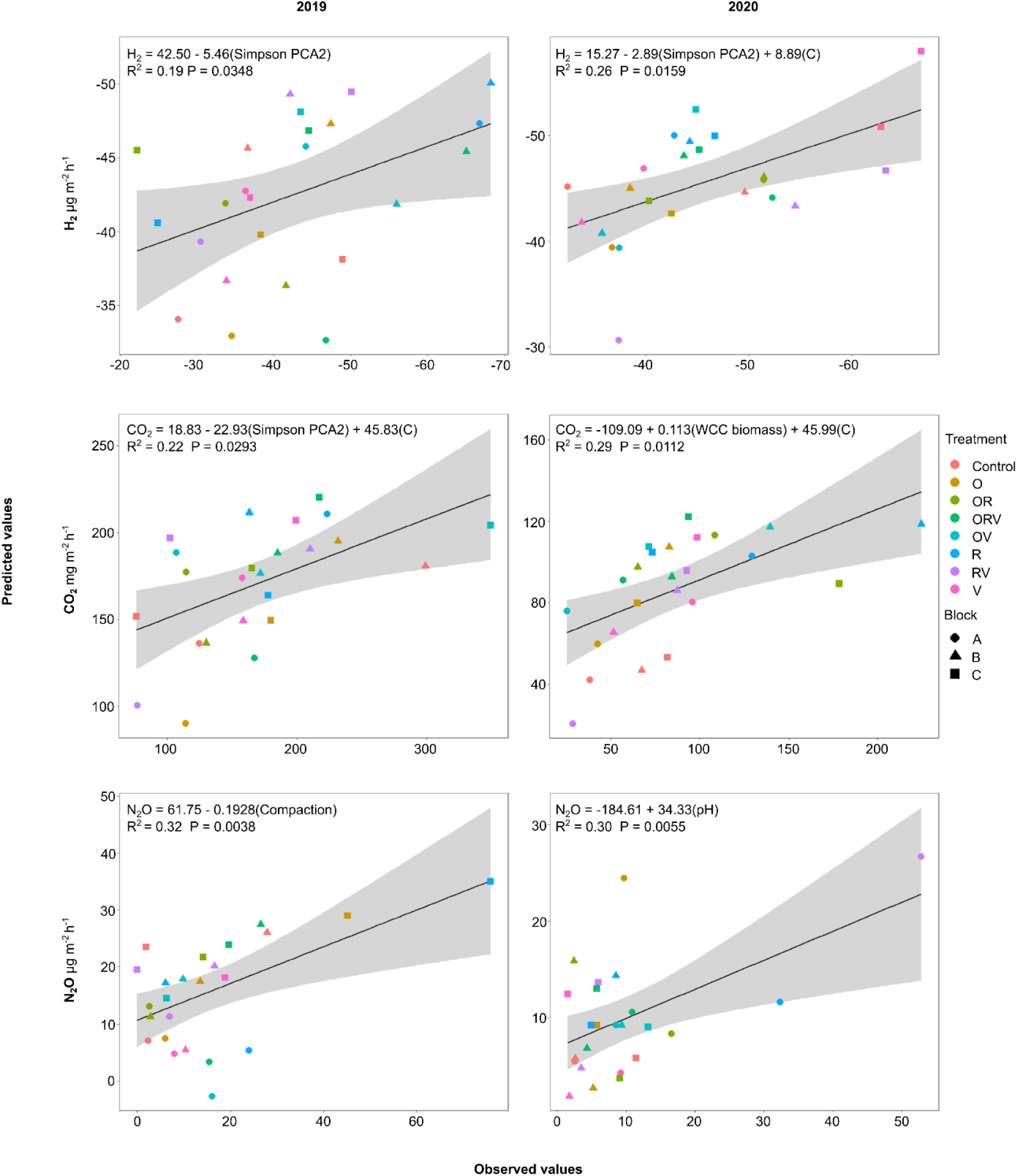
Predicted gas fluxes values for each plot plotted against observed gas fluxes median values. Equations of the predicted gas values are indicated on each facet with their *P*-value and R^2^ coefficient for the single or multiple linear regressions. R^2^ values refer to the adjusted R^2^ for the multiple linear regressions.

## Discussion

Despite the predictable benefits of WCC on carbon storage and soil protection against erosion, other benefits including enhanced microbial diversity and delivery of ecosystem services related to nutrient cycling and pest control appear more variable among different sites. Beside difficulties in predicting the fate of microbial diversity to WCC, there is no threshold to define optimal diversity targets. This is partly explained by shaded relationships between microbial diversity and ecosystem functioning by functional redundancy in microbes and the impact of microbial interaction in supporting functions. Here, the relationship between aboveground vegetation biomass diversity and soil microbial diversity and functioning related to trace gas exchanges was examined in a short-term WCC trial. The legacy of WCC treatment on soil chemical and biological features was decoupled from trace gases fluxes. Sensitivity of trace gas fluxes to local variations of soil features was related to the diversity of microbial communities involved in processes, with the guild of HOB displaying the lowest species richness appearing the most responsive.

### Legacy effect of WCC treatments

While the long-term legacy effects of WCC on nitrogen availability and SOC accumulation have been examined, there is a lack of information regarding the delay before experiencing a WCC legacy effect in agroecosystems. In this study, neither soil microbial diversity nor trace gas fluxes responded to the various combinations of WCC treatment encompassing one, two or three species. A delayed legacy effect of WCC treatments was noticed on fungal communities. Their composition was more homogeneous and less diverse in the hairy vetch treatment compared to the mixture of vetch and radish, in the second cash crop rotation sequence (figure S2). These results suggest that hairy vetch alone can limit fungal community diversity at the landscape level but the exact mechanisms responsible for the delay remain obscure. There is no consensus on the time needed before WCC drives noticeable compositional change in soil fungal communities. Njeru et al. (2014) found a delayed legacy effect with a reduced diversity of arbuscular mycorrhizal in vetch alone when compared to WCC mixtures. In contrast, compositional changes of fungal communities were observed in less than one year after WCC integration in crop rotation sequences (Cloutier et al., 2020). Different aspects including the sampling time following green manure integration in soil, the biomass yield of WCC and soil management practices are drivers of the response of fungal communities to WCC treatments. The observation that fungal communities are more responsive than bacteria to WCC treatment is in line with a previous investigation (Castle et al., 2021). In this study, neither bacterial, fungal, HOB or denitrifying community structure and abundance displayed a variation pattern related to WCC treatment. Despite the absence of WCC treatment effect on trace gas fluxes measured in the field trial, the combination of WCC biomass with soil microbiological and physicochemical features contributed to explain variations in trace gas fluxes.

### Gas-specific response to WCC biomass and soil features

In general, measured fluxes were comparable to previous reports from agroecosystems. CO_2_ net emission ranging between 200 and 2000 mg CO_2_ m^-2^ h^-1^ were observed in clayey soil after cover crop application (Hendrix et al., 1988; Kallenbach et al., 2010; Maljanen et al., 2001). CO_2_ flux reported in this study are within or below this range, with values comprised between 0 and 743 mg CO_2_ m^-2^ h^-1^ (table 1). Measured N_2_O fluxes between -130 and 634 µg N_2_O m^-2^ h^-1^ were comparable to those reported from clayey soil after cover crop application, -2 to 1200 µg N_2_O m^-2^ h^-1^ (Guardia et al., 2016; Maljanen et al., 2001). H_2_ flux measurements from agroecosystem are scarce, but an annual average of -14 µg H_2_ m^-2^ h^-1^ was observed in a temperate grassland using a micrometeorological technique (Constant et al., 2008) which is in the same magnitude than field observations -2 and -243 µg H_2_ m^-2^ h^-1^ (table 1). Although the spatial variation of physicochemical and microbiological soil attributes was constrained due to the scale of the field trial relative to multiple sites analyses, a number of parameters were identified to explain variation of measured trace gas fluxes.

Soil parameters explaining variation of CO_2_ and N_2_O fluxes were inconsistent among both growing seasons. Soil total carbon was the most important and consistent variable in explaining CO_2_ fluxes due to the promotion of microbial respiration in response to SOM content in soil (Alvarez & Alvarez, 2000). Spatial variation and the incorporation of green manure are two potential drivers of soil carbon content variation in 2019, whereas no supplementary gain attributed to WCC biomass can be attributed in the growing season of 2020. A significant relationship was observed between WCC biomass and CO_2_ fluxes in 2020 only. The lack of such a relationship in 2019, when WCC residues and living biomass were incorporated in soil is likely explained by the incidence of other limiting factors of CO_2_ emissions including, for instance, lower rain precipitation. Nevertheless, the fact that CO_2_ fluxes measured in the different WCC treatments were not distinguishable implies that cover crop species are less important than the C input in soil as biomass in driving CO_2_ emissions (Bavin et al., 2009; Schlesinger, 1984). Variation of soil microbial diversity within the field trial was not sufficient to explain the pattern of CO_2_ fluxes. The absence of a significant relationship between WCC biomass and N_2_O emissions is in agreement with the result of a meta-analysis reporting the absence of N_2_O flux promotion in 40% analyzed studies (Basche et al., 2014). The highest N_2_O emissions were observed in June (figure 3) following fertilizer application. Indeed, the N input of fertilisers supplies microbial nitrification and denitrification processes, leading to enhanced N_2_O emissions (Ruser et al., 1998). Bavin et al. (2009) showed that N fertilisation and fertilizer type are the dominant factors controlling N_2_O fluxes in maize-soybean rotation system including cover crop. In this study, a negative relationship was noticed between N_2_O emissions and soil compaction in 2019, whereas a positive correlation with soil pH was noticed in 2020. The significance of these relationships is obscured by the small range of pH and compaction values measured among the plots of the experimental trial. Neither the composition of microbial communities nor their alpha diversity profile was related to variation of N_2_O fluxes. The broad diversity of microorganisms contributing to CO_2_ and N_2_O fluxes and constrained variation of microbial diversity at the scale of the field trial are two potential explanations for the decoupling between microbial diversity and process rate.

H_2_ fluxes are limited by gas diffusion in soil constrained by soil porosity. Forest ecosystems display higher H_2_ uptake rates than agroecosystems (Ehhalt & Rohrer, 2009). The exact mechanisms steering these different performances are unknown, but a combination of biological and physicochemical attributes is expected to contribute to the biological sink of atmospheric H_2_. Until now, most of the investigations of environmental factors driving H_2_ oxidation activity were examined in soil microcosms where soil water content and texture deciphering gas diffusion limitations are well constrained. Khdhiri et al. (2015) showed that 92 % of the H_2_ oxidation rate variance can be explained by the interaction of soil total carbon content and the HOB relative abundance in soil. Other factors including soil pH, nitrate concentrations, substrate-inducted respiration and soil potential nitrification were related to H_2_ oxidation activity in soil (Gödde et al., 2000). With the exception of total carbon content in 2020, neither physicochemical nor diversity variables were related to H_2_ soil uptake rate measured in the field trial. This is in part explained by the small variability of soil properties among the plots of the field trial examined in this study. Here, the soil uptake of atmospheric H_2_ displayed a spatial distribution pattern in the experimental field trial, which varied between the two years. The pattern was better explained by alpha than beta diversity of microbial communities. HOB functional gene or bacterial 16S rRNA gene alone were insufficient to capture flux variation. The alpha diversity composite variable based on the Simpson index composite metric PCA2 was the only variable explaining H_2_ fluxes variation for both years. The relationship between ecosystem functioning and the evenness of microbial communities has received little attention in previous reports (Bender et al., 2016; Le Bagousse-Pinguet et al., 2021; Nielsen et al., 2011; Wittebolle et al., 2009). Beside the beneficial impact of species richness ensuring representation of functions, alpha diversity can influence their expression through various ecological filtering effects, such as microbial interaction (Ho et al., 2014). For instance, the proliferation and activity of microorganisms are expected to be driven by a net balance of competition and complementarity with the other members of the community. In studies integrating multiple land-use types of sites encompassing gradients of climate or environmental features, the interplay of competition and complementarity is often best represented by compositional change of microbial communities. At the scale of the field trial, composition of microbial communities among the plots was relatively constrained, supporting a most important contribution of evenness in defining competition and complementarity boundaries of HOB responsible for H_2_ uptake activity in soil. This importance of evenness was elegantly supported in 100 synthetic communities with the same richness, but different evenness. Examination of microbial successions among microbial assemblies led to the identification of species favored in synthetic communities characterized by elevated or low evenness (Ehsani et al., 2018). H_2_ soil uptake rate was the sole process related to spatial variation of microbial communities. This is likely explained by their lower diversity than the other guilds mediating N_2_O and CO_2_ emissions. The biological sink of atmospheric H_2_ is generally assumed limited by gas diffusion in soil, but the observations form this WCC trial supports the importance of soil microbial communities as well.

In conclusion, fungal communities were the most sensitive to integration of WCC in the crop rotation. At the scale of this short-term field trial, the integration of WCC in crop rotation was not sufficient to promote belowground microbial diversity expressed by examining bacteria, fungi, HOB and denitrifying bacteria. Similarly, WCC were not sufficient to reduce N_2_O emissions and promote the biological sink of H_2_ in summertime. The variation of CO_2_, N_2_O and H_2_ flux explained by different factors supports the importance of examining multiple gases to evaluate benefits of agroecosystem management practices on trace gas exchanges.

## Supporting information

Supplemental method and result

## Acknowledgements

This work was supported by the “*Programme innov’action agroalimentaire*” through the Quebec-Ontario cooperation for agri-food research competition (project number IA118802). The work of XB was supported by a Natural Sciences and Engineering Research Council of Canada (NSERC) M.Sc. scholarship in 2020 and a “*Fonds de recherche du Québec – Nature et technologies*” M.Sc. scholarship in 2021.

## Notes

### Competing Interest Statement

The authors have declared no competing interest.

