## Supplemental method and result for "The biological sink of atmospheric H2 is more sensitive to spatial variation of microbial diversity than N_2_O and CO_2_ emissions in an agroecosystem"

**Supplementary method 1**

*Droplet digital PCR assays*

Dilution of the DNA template were effectuated and tested while varying the annealing temperature from 51 to 63 ºC to determine the appropriated parameter for each gene tested. Primer final concentration of 100 nM was used for all assays. Each sample reaction mix had a total volume of 21.4 µl and was composed of 11.25 µl of Evagreen Supermix (Bio-Rad, cat no. 1864043), 1.125 µl of both forward and reverse primers (2 µM), 2.9 µl of sterilised Milli-Q water and 5 µl of DNA template. The samples solutions were prepared in a 96-well Semi-Skirted ddPCR plate (Bio-Rad, cat no. 12001925). 20 µl of each sample was loaded in the Droplet Generator DG8™ Cartridge (Bio-Rad, cat no. 1864008) afterwards 65 µl of QX200 Droplet Generation Oil for EvaGreen (Bio-Rad, cat no. 1864005) were added in the proper well. The droplet emulsion was then pipetted in a clean 96-well plate which was sealed at 180 °C in the PX1™ PCR Plate Sealer (Bio-Rad) using a pierceable foil heat seal (Bio-Rad, cat no. 1814040). For each assay, 50 PCR cycles of the cycling protocol shown in table S1 were performed.

Table S1. Cycling condition for Bio-Rad's C1000 Touch Thermal Cycler used for the assays.

| Temperature (°C) | Time | Number of Cycles |
|---|---|---|
| 95 | 5 min | 1 |
| 95 | 30 sec | 50 |
| Varying | 1 min | |
| 72 | 30 sec | |
| 4 | 5 min | 1 |
| 90 | 5 min | 1 |
| 12 | Infinite | 1 |

* ramprade of 2 °C/s, heated lid set to 105 °C and sample volume set to 40 µl.

Table S2. ddPCR assays parameters.

| ddPCR assays | DNA templates concentration | Annealing temperature (°C) | Manual threshold setting |
|---|---|---|---|
| 16S | $2 \cdot 10^{-4}$ | 51.0 | 4800-8700 |
| ITS | $4 \cdot 10^{-3}$ | 53.4 | 9250 |
| *hhyL* | $2 \cdot 10^{-2}$ | 63.0 | 6100-8000 |
| *nosZ* | $2 \cdot 10^{-2}$ | 62.3 | 7150-9300 |

* ramprade of 2 °C/s, heated lid set to 105 °C and sample volume set to 40 µl. DNA template concentration refer to the dilution factor of the extracted DNA samples.

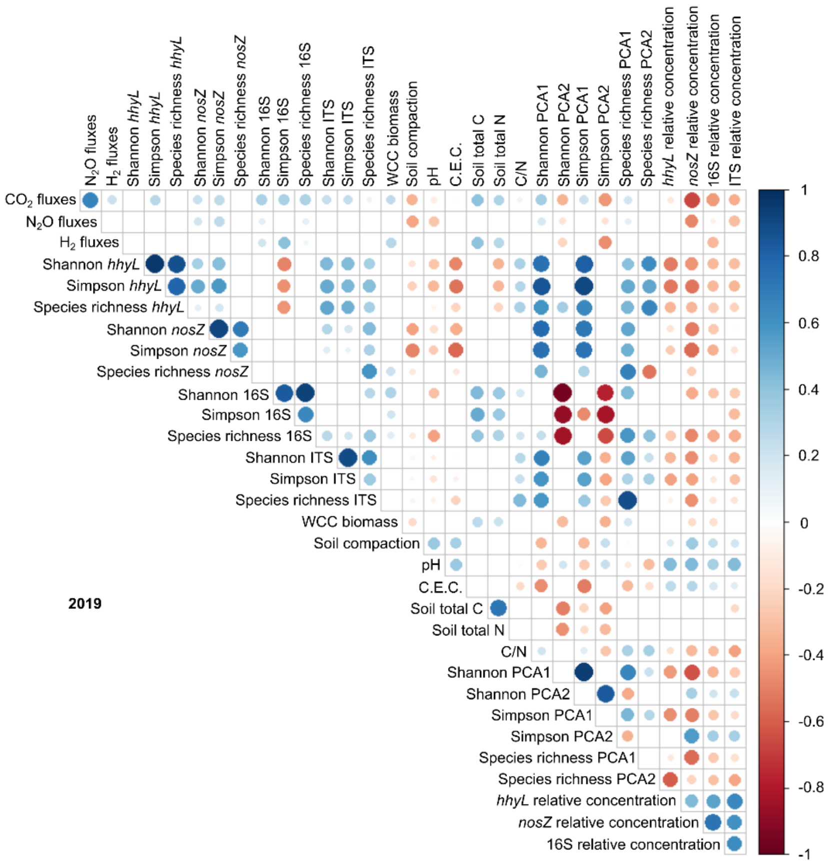
N2O fluxes
H2 fluxes
Shannon hhyL
Simpson hhyL
Species richness hhyL
Shannon nosZ
Simpson nosZ
Species richness nosZ
Shannon 16S
Simpson 16S
Species richness 16S
Shannon ITS
Simpson ITS
Species richness ITS
WCC biomass
Soil compaction
pH
C.E.C.
Soil total C
Soil total N
C/N
Shannon PCA1
Shannon PCA2
Simpson PCA1
Simpson PCA2
Species richness PCA1
Species richness PCA2
hhyL relative concentration
nosZ relative concentration
16S relative concentration
ITS relative concentration
CO2 fluxes
1
0.8
0.6
0.4
0.2
0
-0.2
-0.4
-0.6
-0.8
-1
2019

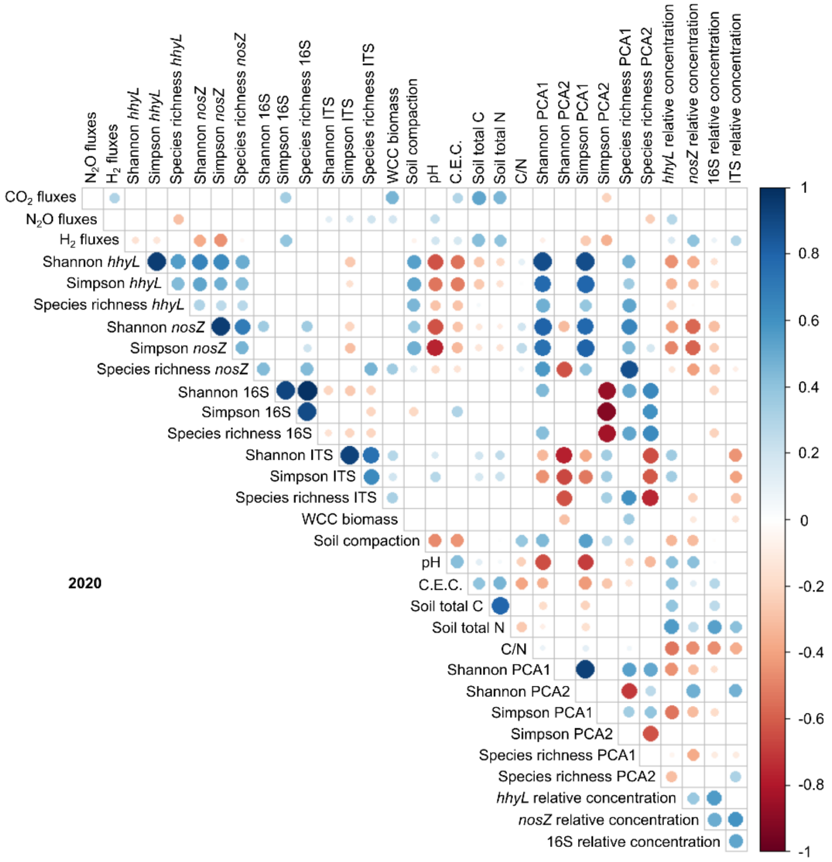
N2O fluxes
H2 fluxes
Shannon hhyL
Simpson hhyL
Species richness hhyL
Shannon nosZ
Simpson nosZ
Species richness nosZ
Shannon 16S
Simpson 16S
Species richness 16S
Shannon ITS
Simpson ITS
Species richness ITS
WCC biomass
Soil compaction
pH
C.E.C.
Soil total C
Soil total N
C/N
Shannon PCA1
Shannon PCA2
Simpson PCA1
Simpson PCA2
Species richness PCA1
Species richness PCA2
hhyL relative concentration
nosZ relative concentration
16S relative concentration
ITS relative concentration
CO2 fluxes
1
0.8
0.6
0.4
0.2
0
-0.2
-0.4
-0.6
-0.8
-1
2020

**Figure S1.** Spearman correlation matrix between gas fluxes, soil physicochemical proprieties, diversity index and genes copy number relative concentration for 2019 and 2020. The presence dot represents a significant correlation ($P < 0.05$). Dot size is proportional to the rho correlation coefficient value ($\rho$), while the red color is associated with a negative correlation, and the blue with a positive correlation.

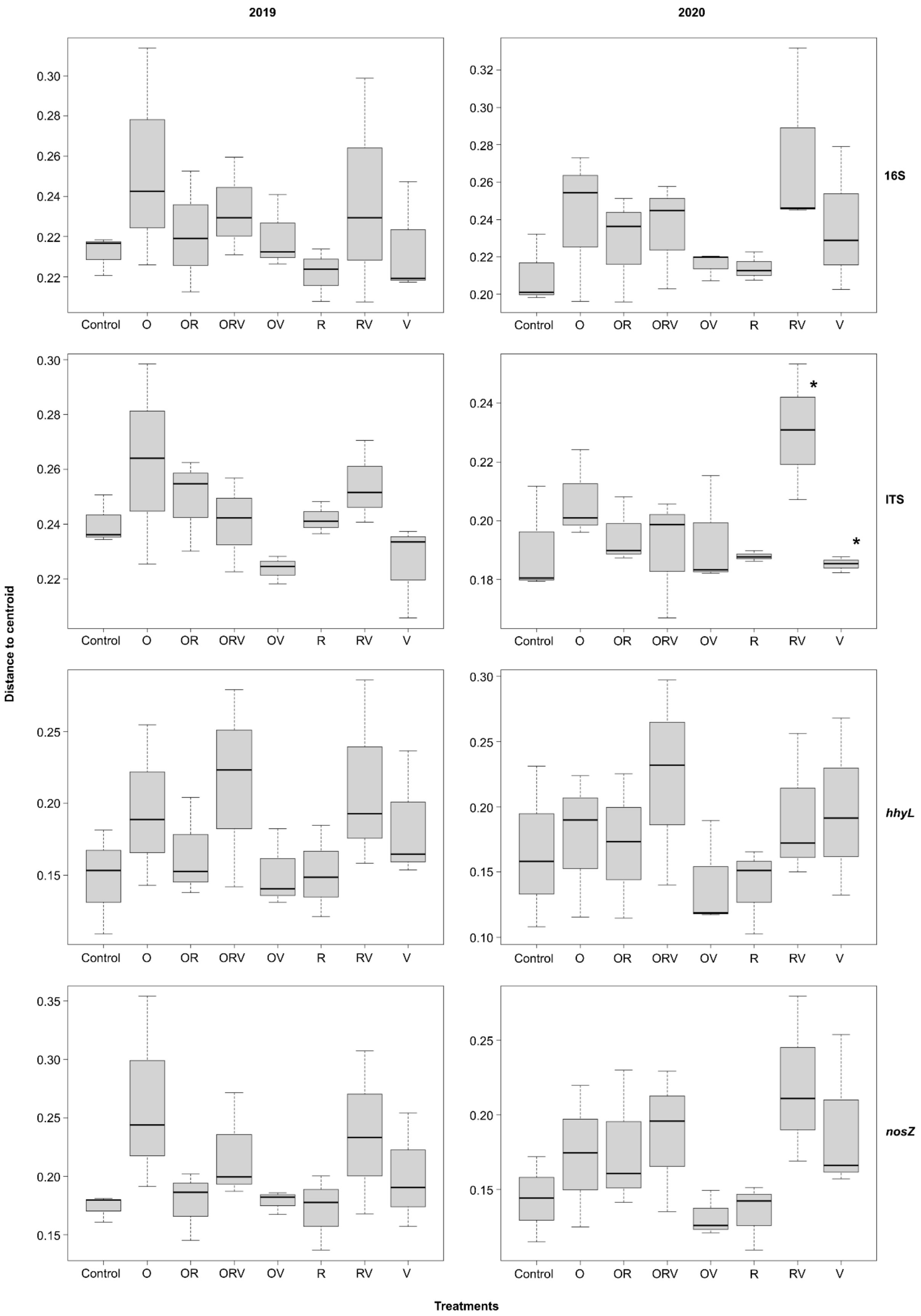

2019
2020
16S
ITS
hhyL
nosZ
Distance to centroid
Treatments
Control
O
OR
ORV
OV
R
RV
V
*

**Figure S2.** Dispersion of microbial community structures from replicated WCC treatments for four genes in 2019 and 2020, respectively. Treatments with significantly different distances to the centroid are represented by *.

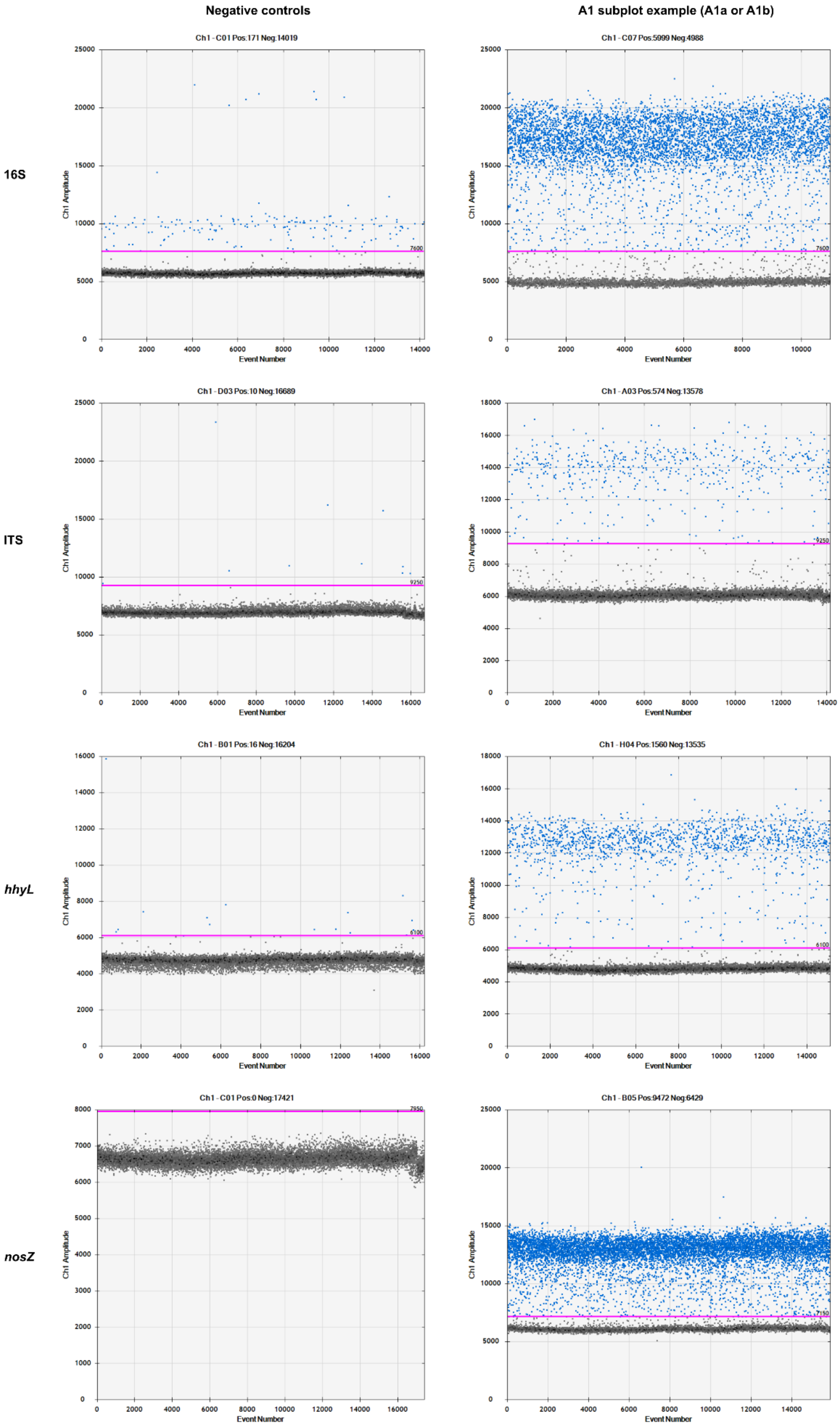

Negative controls
A1 subplot example (A1a or A1b)
16S
ITS
hhyL
nosZ
Ch1 - C01 Pos:171 Neg:14019
Ch1 - C07 Pos:5999 Neg:4988
Ch1 - D03 Pos:10 Neg:16689
Ch1 - A03 Pos:574 Neg:13578
Ch1 - B01 Pos:16 Neg:16204
Ch1 - H04 Pos:1560 Neg:13535
Ch1 - C01 Pos:0 Neg:17421
Ch1 - B05 Pos:9472 Neg:6429
Ch1 Amplitude
Event Number

**Figure S3.** ddPCR graphical display of 1D amplitude of the first negative control and the A1a or A1b subplot with their threshold setting for each ddPCR assays on the 2019 samples

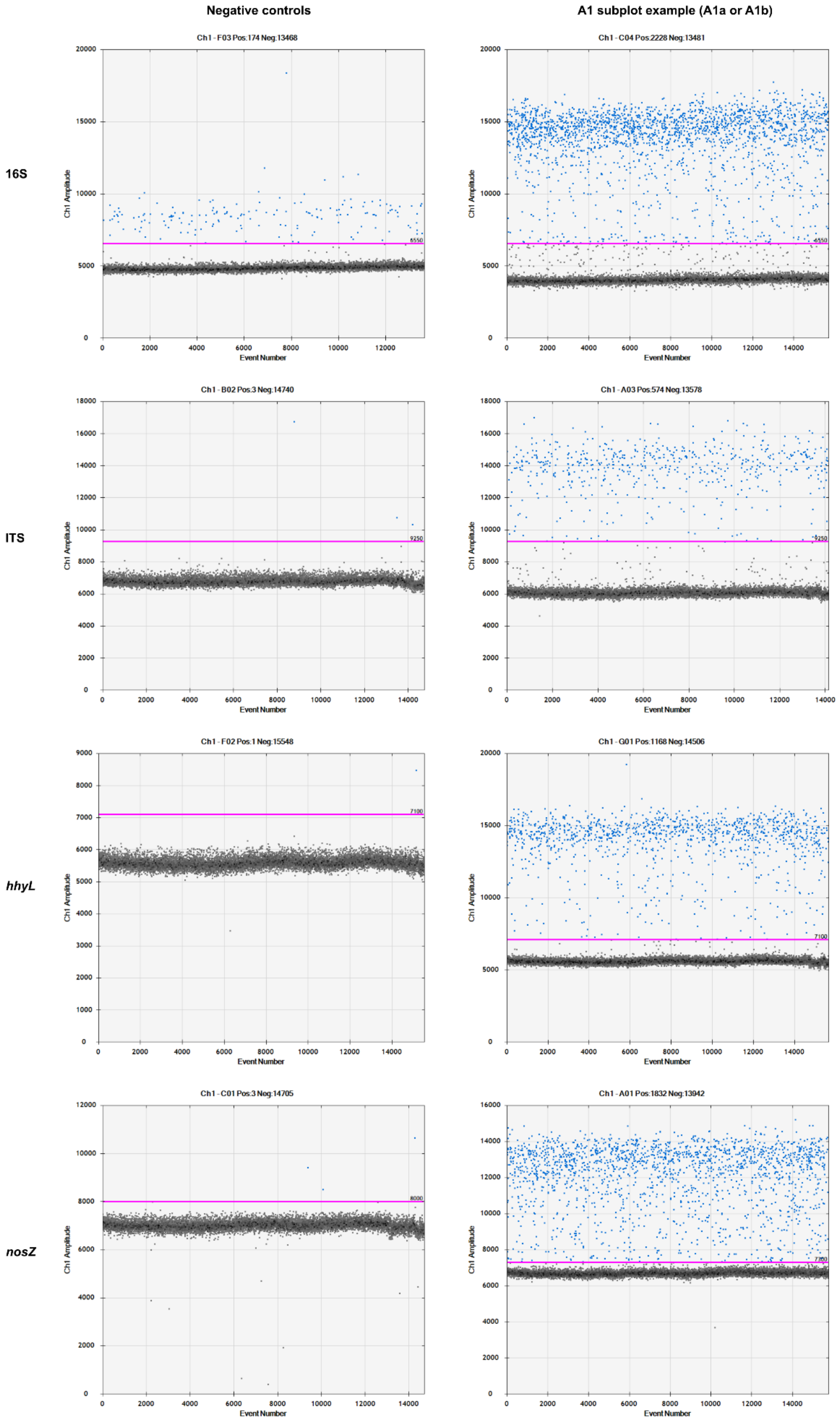
Negative controls
A1 subplot example (A1a or A1b)
16S
ITS
hhyL
nosZ
Ch1 - F03 Pos:174 Neg:13468
Ch1 - C04 Pos:2228 Neg:13481
Ch1 - B02 Pos:3 Neg:14740
Ch1 - A03 Pos:574 Neg:13578
Ch1 - F02 Pos:1 Neg:15548
Ch1 - G01 Pos:1168 Neg:14506
Ch1 - C01 Pos:3 Neg:14705
Ch1 - A01 Pos:1832 Neg:13942
Ch1 Amplitude
Event Number
6550
9250
7100
8000
7300

**Figure S4.** ddPCR graphical display of 1D amplitude of the first negative control and the A1a or A1b subplot with their threshold setting for each ddPCR assays on the 2020 samples
